# Fiber-TEnCATS reveals haplotype-specific chromatin accessibility and DNA methylation at human L1HS loci

**DOI:** 10.64898/2026.06.26.734832

**Authors:** Katarina Pavlovic, Torrin L. McDonald, Adam G. Diehl, Jessica A. Switzenberg, Alan P. Boyle

**Author notes:** To whom correspondence should be addressed. (A.P.B.).

## Abstract

Human-specific long interspersed nuclear element-1 (L1HS) is an active and autonomous retrotransposon in the human genome. Changes in its transcription and transposition are known to affect cellular processes involved in development and aging, and diseases such as neurological disorders and cancer. To better understand natural variability in epigenetic patterns that affect L1HS regulation, we developed a targeted long-read method to simultaneously profile individual haplotypes for DNA methylation and chromatin accessibility across L1HS loci in a healthy human cell line trio. We show that the intronic L1HS in the ZNF638 gene consistently displays high chromatin accessibility and DNA hypomethylation with bidirectional transcription. Our approach also reveals additional intronic and intergenic L1HS copies with allele-specific chromatin accessibility and methylation, and instances of reduced promoter DNA methylation that does not correspond with increased chromatin accessibility. We also identify potential cases of non-Mendelian inheritance of DNA methylation patterns over a subset of L1HS promoters. Our method’s high coverage over L1HS loci enables detection and profiling of loci that are missed even by long-read-based assemblies and enables more accurate inheritance tracing of L1HS insertions. Overall, our results offer new insights into the locus-specific regulation of both reference and non-reference L1HS within the human genome.

## Introduction

Long interspersed nuclear element-1 (L1) retrotransposons are the second most abundant and highly polymorphic family of transposable elements (TEs) in humans^1–3^. They are also the only family of TEs with members that can autonomously mobilize in the modern human genome^4^. The evolutionarily youngest and still transpositionally active member of this family, L1HS, creates a new insertion approximately every 60 live births^5^.

L1HS has a bidirectional promoter. An accessible L1HS sense promoter is crucial for the transcription and subsequent mobilization of the element^6,7^, while an antisense promoter can initiate transcription into the flanking genomic sequence^8,9^. Transcription from the antisense promoter can influence both the transcription and translation of genes that L1HS inserts into or nearby, forming spliced chimeric transcripts between L1HS and gene exons and, in some cases, fusion proteins^8–10^. Two L1HS proteins, ORF1p and ORF2p, are necessary not only for the element’s own transposition, but also for the transposition of other nonautonomous families of transposable elements, such as Alus and SVAs^11–13^. During transposition, ORF1p acts as a nucleic acid chaperone^14^ while ORF2p has reverse transcriptase and endonuclease activities^15^.

L1HS elements that become transcriptionally active and retrotransposition-competent can influence genome integrity and function^16–19^. As a result, host cells have evolved mechanisms to try to limit L1HS transcription. While KZFP/TRIM28 complexes effectively silence several ancestral L1 lineages, the L1HS subfamily has accumulated mutations in its promoter that enable these TEs to bypass KZFPs important for TRIM28 recruitment^20,21^. Consequently, DNA methylation of the CpG-rich promoter is believed to be the primary mechanism for silencing L1HS at the transcriptional level^8,22,23^. In most differentiated somatic cells, extensive methylation across L1HS promoters largely prevents transcription. However, while promoter hypomethylation can lead to reactivation, it does not always guarantee it^6^, suggesting that additional features of the local chromatin environment also contribute to L1HS silencing. For instance, L1HS insertions within heterochromatic regions can acquire local repressive histone marks that promote a less accessible chromatin state, limiting transcription factor binding and transcriptional activation regardless of the promoter’s methylation status^7^. Even in euchromatic regions, L1HS insertions, especially those located within introns of active genes, are often targeted by the HUSH complex, which deposits repressive histone marks^24^. Additionally, L1HS loci with hypomethylated promoters are detected by the HUSH-MORC2 pathway, which prevents their reactivation by adding H3K9me3 marks and reinforcing CpG methylation^25,26^.

Instead of assaying individual histone modifications, chromatin accessibility can be used as a broader readout of whether the local chromatin environment permits transcription factor binding and transcriptional initiation. Most efforts to study L1HS chromatin accessibility beyond DNA methylation rely on short-read sequencing^7,27^. Due to the high sequence similarity among L1HS loci and their length, which often exceeds the read size, short-read methods usually struggle to capture signals along the entire length of the element, beyond the promoter region immediately adjacent to the unique flanking genomic sequence. This limits the ability to study chromatin accessibility across the rest of the L1HS sequence at different genomic loci. Limited length of short reads also reduces the probability that any individual read will overlap one or more heterozygous variants and be successfully assigned to its haplotype of origin.

Long-read sequencing technologies now enable us to better study the complex regulation of L1HS elements. We can unambiguously haplotag and map to specific genomic L1HS loci, as reads can cover the entire L1HS element and provide sufficient flanking unique genomic sequences ^28,29^. These technologies also enable direct measurement of DNA methylation^30^. And more recently, methods have been developed that combine long-read sequencing with methyltransferases to enable simultaneous measurement of chromatin accessibility^31–33^.

Here, we employ a new TE-optimized long-read method, Fiber-TEnCATS (**T**ransposable **E**lement **n**anopore **Ca**s9-**T**argeted **S**equencing with **Fiber**-seq^31^ chromatin accessibility information), to simultaneously analyze haplotype-specific sequences, DNA methylation, and chromatin accessibility across the full length of both reference and non-reference L1HS, and apply it to the healthy human cell line trio HG002, HG003, and HG004. This design enables us to investigate how important contributors to L1HS silencing differ between loci within one individual and between related individuals, and to comprehensively uncover their variability across the full length of different L1HS genomic copies.

## Results

### Fiber-TEnCATS Enables High-Resolution Profiling of L1HS Elements

Fiber-TEnCATS is an extension of our previously published TEnCATS that uses a single guide RNA (sgRNA), recognizing a subsequence unique to L1HS consensus, to direct Cas9 to target and cut within genomic L1HS loci before ONT sequencing^29^. However, prior to the TEnCATS protocol, regions of accessible chromatin are labeled with an exogenous m6A mark, selectively introduced by Hia5 methyltransferase^31^. Nanopore sequencing then detects both the standard DNA bases and their modifications, enabling the simultaneous profiling of endogenous DNA methylation (methyl-CpG) and exogenous m6A as an indicator of chromatin accessibility (Figure 1A). Using a single PromethION flow cell, we captured over 96% of genomic L1HS loci with a PAM site and up to 3 mismatches relative to the sgRNA (Figure 1B). In this analysis, we included only L1HS loci with at least eight reads of coverage, which we considered the minimum for chromatin accessibility analysis. On average, our method provided more than twice the coverage of L1HS regions compared to the standard Fiber-seq method^31^ (Figure 1C), reducing the number of flow cells needed to reach high sequencing depth and lowering costs. It also recapitulated known ATAC-seq patterns in chromatin accessibility—such as those observed in ENCODE GM12878 data (Figure 1D)—and detected signals deeper within repetitive regions missed by ATAC-seq due to short-read multimapping issues (Figure 1E).

**Figure 1:**
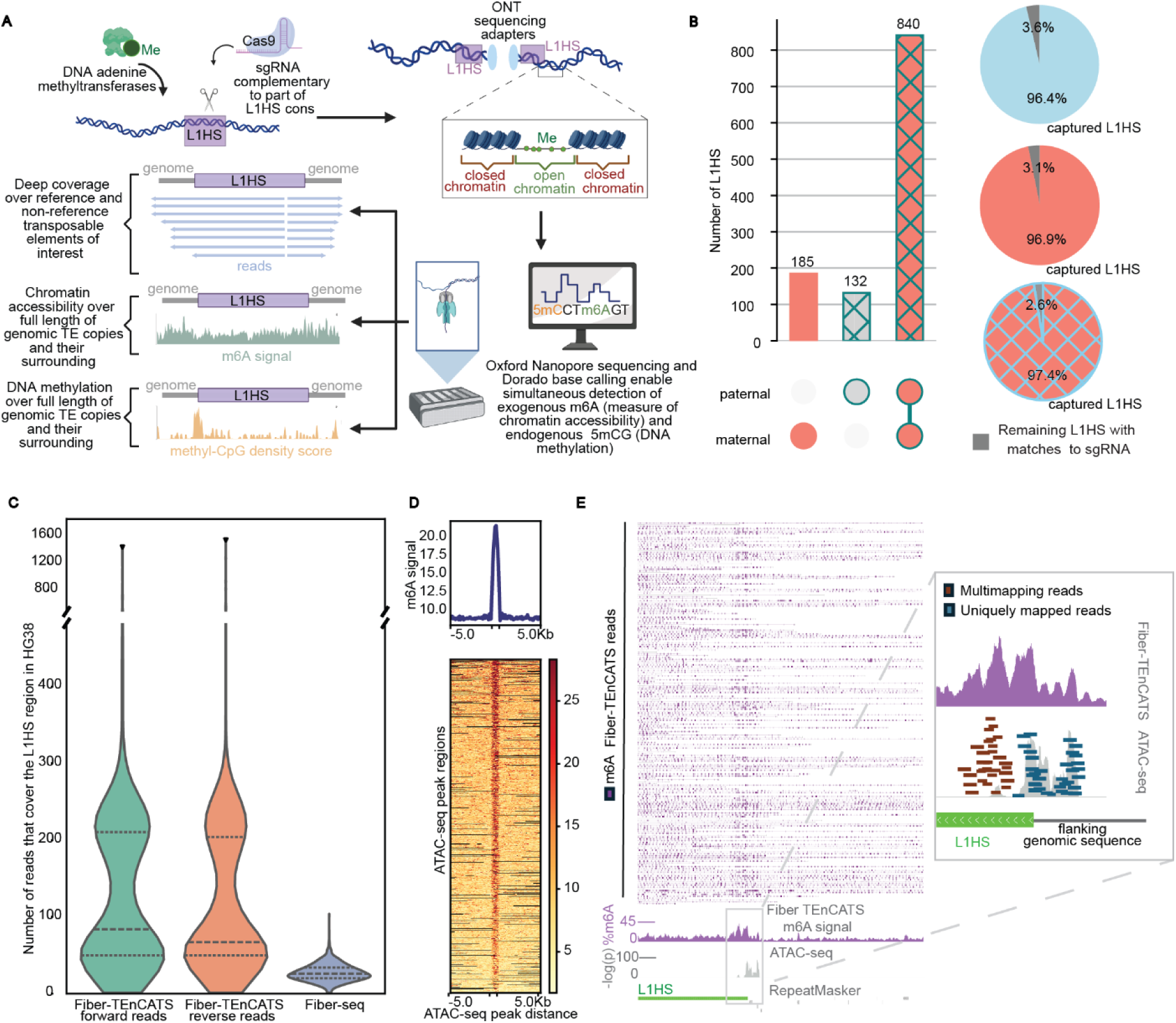
Fiber-seq and targeted enrichment capture of transposable elements – Fiber-TEnCATS. **A)** sgRNA directs Cas9 to genomic L1HS elements, leading to bidirectional sequencing spreading from the adapters placed at the cut site. Hia5 methyltransferase introduces an exogenous m6A mark in accessible chromatin regions. Nanopore sequencing is used to obtain information on both the regular DNA bases and their modifications, including endogenous 5mCpG as a marker of DNA methylation, and exogenous m6A as a marker of accessible chromatin. This section of the panel was created with BioRender.com**. B)** Upset plot shows the number of heterozygous and homozygous L1HS loci from HG002 T2T captured with our method with at least 8 supporting reads. The accompanying pie charts show that captured L1HSs comprise over 96% of capturable loci (with up to 3 mismatches to sgRNA and with a PAM site) from each category in the upset plot. **C)** Fiber-TEnCATS, separated by read strand orientation, shows more than twice the average coverage over L1HS regions compared to the standard Fiber-seq method. **D)** Metagene plot showing enrichment of m6A signal over ATAC-seq peak regions covered by Fiber-TEnCATS reads. **E)** Genome browser in a Box (GBiB) view of Fiber-TEnCATS reads over reference L1HS on the antisense strand with an open promoter. ATAC-seq peak does not cover the full length of the open chromatin region due to multimapping of reads that are fully contained within repetitive sequences. Fiber-TEnCATS is able to recover the full length of the open region.

### L1HS in the ZNF638 intron has an active promoter in related individuals

Fiber-TEnCATS identified accessible chromatin at the promoter of an intronic L1HS element within *ZNF638* (*NP220*), a signal that was only partially detected by ATAC-seq (Figure 1E). Previous studies have reported evidence of antisense promoter activity at this locus across diverse cell types and tissues, including healthy lung tissue, neurons, human embryonic stem cells, and HEK293 cells, with some transcripts containing portions of the L1HS 5′ UTR and ZNF638 exons^4,8,10^. ATAC-seq peaks partially overlapping this promoter have also been observed in HEK293 and HeLa cells, and low CpG methylation has been observed in healthy heart tissue^6,30^. Collectively, this evidence suggests that this L1HS locus may not simply escape host-mediated repression sporadically, but that promoter chromatin accessibility could be actively maintained, potentially across generations. To test this hypothesis, we first checked for inheritance of this epiallele in the HG002 trio using personal genome references (Methods). To determine whether a promoter has accessible chromatin, we used a modified Fiber-seq Inferred Regulatory Elements (FIRE) peak-calling approach^34^ (Methods). The ZNF638-overlapping L1HS promoter was enriched for FIRE elements (FDR<0.05) in all trio members and consistently showed an m6A signal above the outlier threshold, indicating accessible chromatin (Figure 2A and Figure 2B). The m6A signal showed a bimodal distribution with peaks over reported sense and antisense promoter regions (Supplementary Figure 1A). To capture both CpG abundance and methylation level when assessing DNA methylation, we defined a methyl-CpG density score as the sum of methylation probabilities across CpG cytosines within a given window, normalized by the total number of cytosines in that window. For promoter-level analysis, we focused on the densest CpG island within the first 450 bases of the promoter region^8^ (Supplementary Figure 1B) and performed an outlier analysis analogous to the m6A signal one (Methods). We found that the CpG island of the ZNF638-overlapping L1HS promoter was hypomethylated across all cell lines (Figure 2A and Figure 2B).

**Figure 2.**
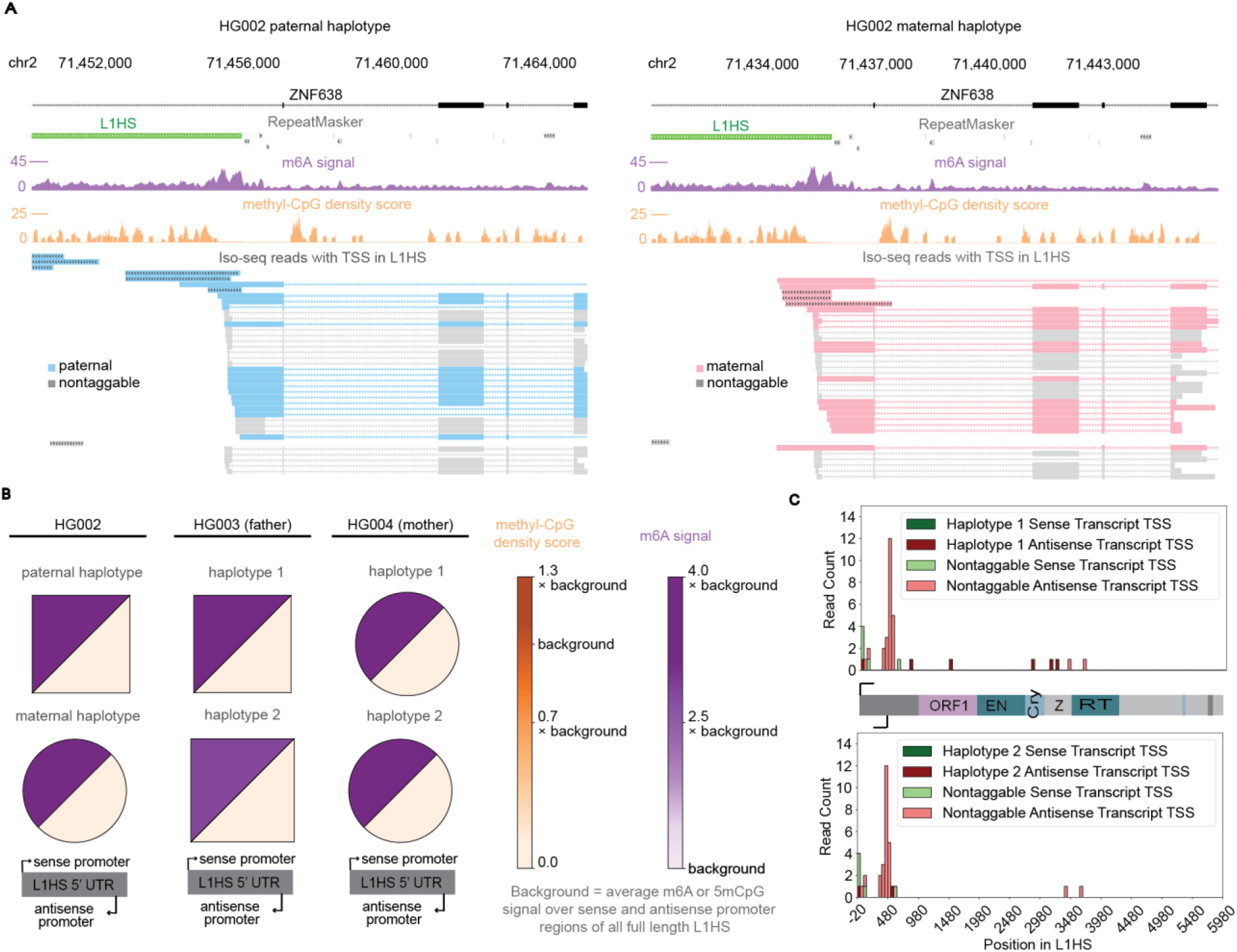
Haplotype resolved view of chromatin accessibility and transcription of L1HS in the ZNF638 intron. **A)** GBiB view of paternal (left) and maternal (right) alleles of L1HS at the ZNF638 locus with corresponding Fiber-TEnCATS and Iso-seq reads. Higher m6A signal at the L1HS promoter is a marker of increased chromatin accessibility in this region on both haplotypes and is accompanied by hypomethylated DNA. Both sense and antisense transcripts are synthesized from this locus in both haplotypes of HG002. Iso-seq reads were filtered to only include transcripts wherein transcription start sites (TSS) or transcription end sites (TES) overlap the given L1HS. **B)** Haplotype-resolved normalized m6A signal and methyl-CpG density score at the ZNF638 L1HS promoter across cell lines. In each tile, the m6A signal is shown in the upper left and methyl-CpG density in the lower right; both were normalized to the average signal across all full-length L1HS promoters in the corresponding sample. Because most full-length L1HS are expected to be silenced in differentiated cells, this background served as an expected signal for the largely silenced L1HS promoter population. **C)** Distribution of TSS of sense and antisense transcripts overlapping ZNF638 L1HS in HG004.

We next assessed methyl-CpG density score patterns across four generations using recently published Platinum Pedigree long-read data from lymphoblastoid cell lines and peripheral blood^35^. The ZNF638 promoter consistently had a low methyl-CpG density score (Supplementary Figure 2A).

Our Fiber-TEnCATS reads were sufficiently long not only to map unambiguously to our region of interest but also to enable read haplotagging^36^, uncovering that all trio samples are homozygous for this epiallele. Using two publicly available Iso-seq datasets from HG002 and HG004, we identified bidirectional transcription in these cell lines (Figure 2A and Figure 2C), further supporting the signatures of accessible chromatin over the promoter. We also found that the majority of antisense transcripts include ZNF638 exons spliced to part of the L1HS 5’ UTR (Figure 2A).

Finally, given its full length and promoter activity, we asked whether this L1HS could be transposition-competent. While the ORF1 sequence appeared intact, ORF2 harbored mutations that introduced a premature stop codon (Supplementary Table 1), making it highly unlikely that this L1HS can transpose autonomously or provide full functional protein machinery for transposition of non-autonomous TEs in the genome.

### Multiple L1HS loci have epiallele-specific promoter accessibility in HG002

Haplotagging Fiber-TEnCATS reads enabled us to observe cases where L1HS epialleles differ in HG002 (Figure 3A). We identified six such cases, four of which were intronic. The L1HS loci in introns of UXS1, XPR1, and TOX genes showed evidence of a potentially active promoter only on the paternal HG002 haplotype. In contrast, the SCFD1 intronic L1HS locus and the 2 remaining intergenic loci had evidence of a potentially active promoter only on the maternal haplotype (Table 1). Across all these loci, methyl-CpG density scores were anticorrelated with the m6A signal at each epiallele (Figure 3B and Supplementary Figure 3). Sequence analysis with L1Xplorer revealed that only XPR1 L1HS locus lacked gaps or frameshift mutations in either of its open reading frames (Supplementary Table 1), making it potentially retrotransposition-competent and a potential source of functional ORF1p and ORF2p for other non-autonomous TEs. To our knowledge, XPR1 L1HS is also the only one of these loci with previously reported retrotransposition assay activity^37^ and recently-transposed progeny^38^.

**Figure 3.**
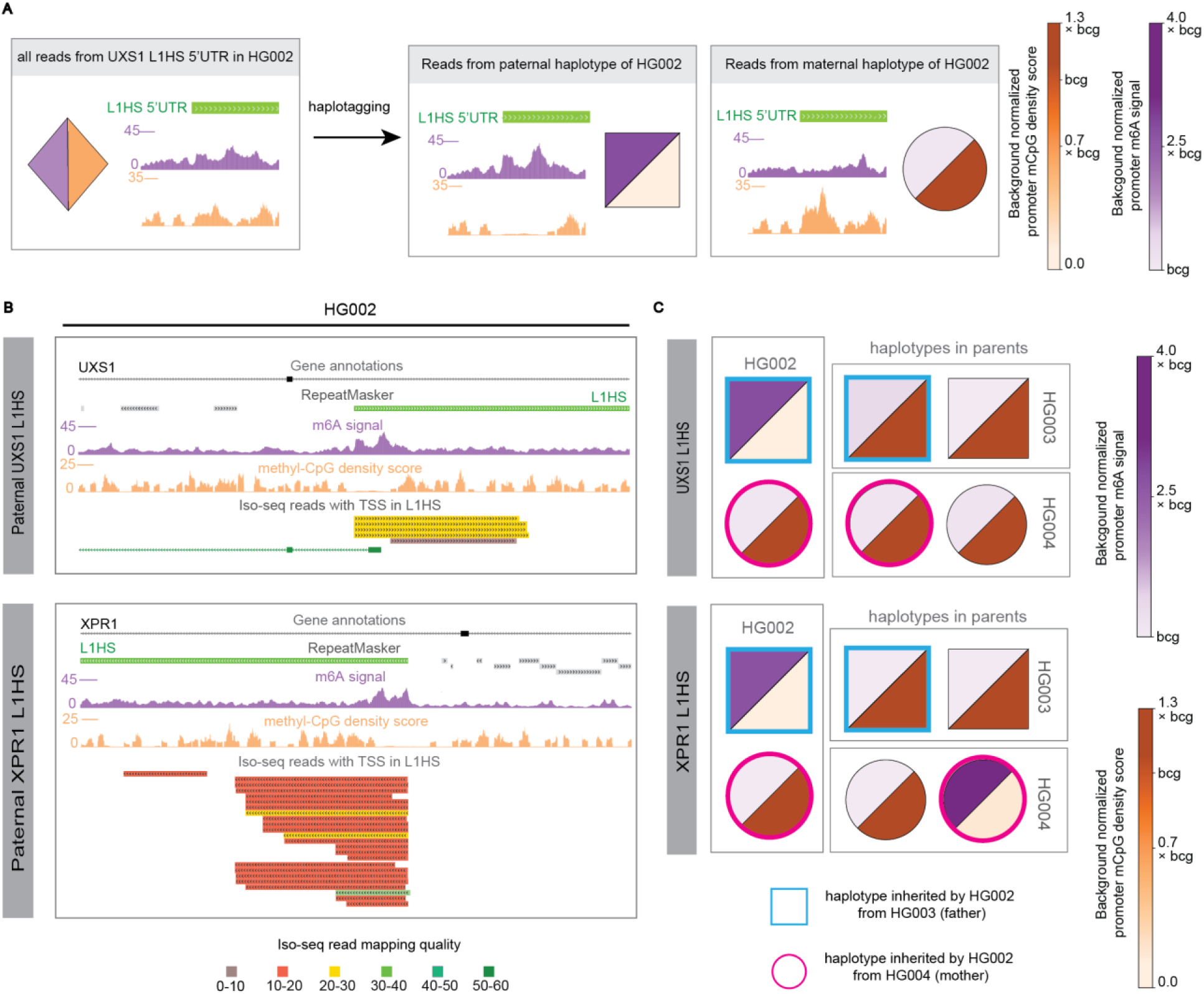
Haplotype-resolved view of UXS1 and XPR1 L1HS loci with heterozygous epialleles in HG002. **A)** Signal over the L1HS promoter in UXS1 intron before and after haplotagging reads – without haplotagging, the m6A peak and low 5mCpG density score in the paternal haplotype would be masked by the opposite pattern in the maternal haplotype and go unnoticed. **B)** GBiB view of L1HS loci in introns of UXS1 and XPR1 genes in the haplotype with an accessible L1HS promoter in HG002. **C)** Haplotype-resolved and normalized DNA methylation (m5CpG) and chromatin accessibility (m6A) signals over promoters of L1HS in UXS1 and XPR1.

**Table 1.**
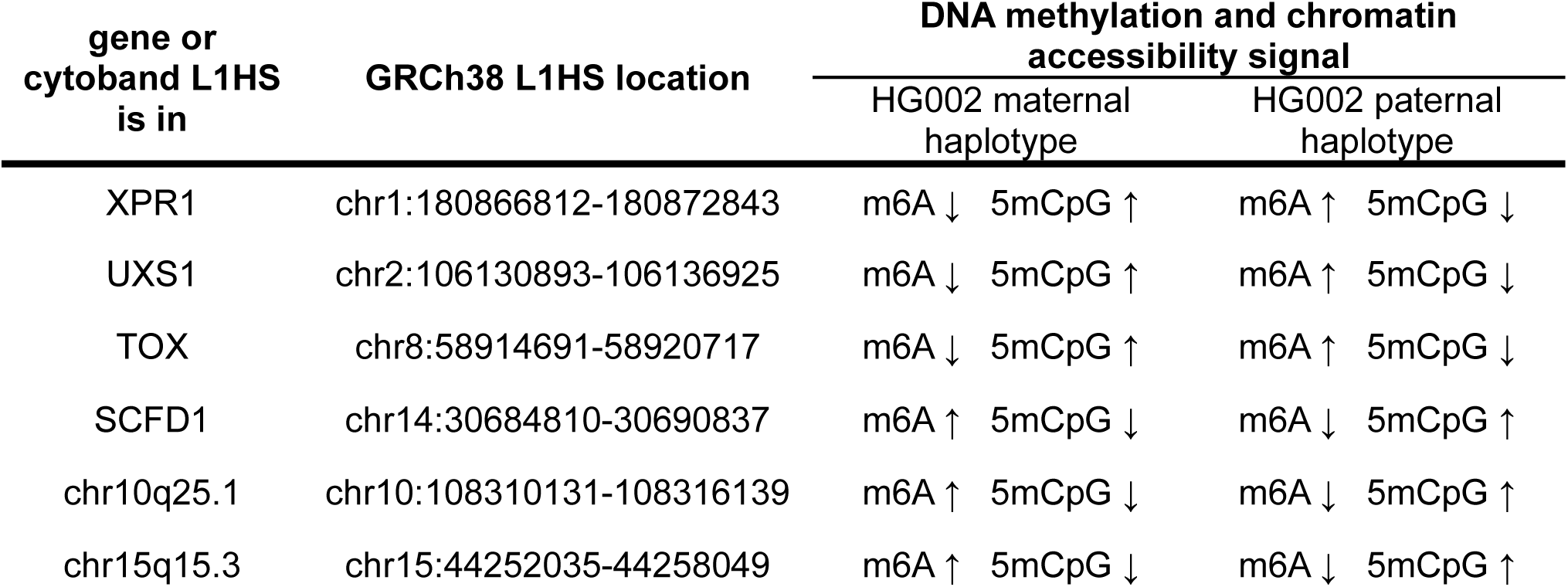
L1HS loci with heterozygous epialleles in HG002 T2T. GRCh38 reference locations of L1HS loci with heterozygous epialleles in the HG002 T2T assembly, along with gene or cytoband labels used in the paper to describe them. For each locus, the table indicates the HG002 haplotype with higher or lower chromatin accessibility and lower or higher DNA methylation. High-accessibility and low-methylation states were defined using outlier thresholds relative to the average across full-length L1HS promoters.

However, using the Wang et al. source inference approach based on internal L1HS sequences and the transduction sequence-based approach (Methods)^39^, we found no evidence that any of these L1HS loci, including XPR1, are recent progenitors of heterozygous L1HS insertions in HG002.

All intronic L1HS loci described above had detectable transcripts, and all except the XPR1 locus showed both sense and antisense transcripts. However, only SCFD1 L1HS transcripts could be haplotagged. Thus, Iso-seq data alone did not reliably identify the transcriptionally active L1HS allele, whereas Fiber-TEnCATS provided allele-resolved context for interpreting the transcript data. Antisense transcription from several of these loci has also been reported, though without haplotype resolution and in different cell types than those in this study. Antisense transcripts of SCFD1^10,40^ and XPR1-overlapping L1HS^4,40^ have been found in embryonic and cancer cell lines, and an ATAC-seq peak partially overlapping the SCFD1 L1HS promoter has been observed in HEK293 cells^6^. Lee et al. reported that expression from the region corresponding to the antisense promoter of the TOX-overlapping L1HS was upregulated in lung cancer compared to normal tissue, but this TE was annotated as L1PA2 in the reference genome used at the time^41^. To our knowledge, no antisense transcripts of UXS1 L1HS or the two intergenic L1HS loci have been reported.

### L1HS promoter methylation shows potential non-Mendelian inheritance without consistent chromatin accessibility changes

Next, we examined chromatin accessibility and DNA methylation at all six epiallele-specific loci in HG003 and HG004 (Figure 3B and Supplementary Figure 3). At half of the heterozygous epiallele loci (UXS1, TOX, and chr10q25.1), all parental haplotypes lacked detectable FIRE enrichment and had hypermethylated promoters. The HG002 maternal allele at the chr15q15.3 L1HS locus showed a 1.6-fold higher normalized m6A signal and a more than 28 percentage points lower normalized methyl-CpG score than the HG004 haplotype being transmitted, whereas the non-inherited allele was consistent with the HG002 maternal allele. The other two L1HS, located in the SCFD1 and XPR1 genes, exhibited heterozygous epialleles in one of HG002’s parents and appeared to be silenced in the other parent. At both loci, the epiallele with a lower promoter methyl-CpG density score and higher m6A signal in HG002 was inherited from a parental haplotype whose L1HS had methyl-CpG and m6A signals characteristic of silencing (Supplementary Figure 4).

As before, we compared chromatin accessibility observations with patterns in promoter methyl-CpG density scores in the Platinum Pedigree^42^ data (Figure 4A and Supplementary Figure 2). Similar to the trio data, the promoter CpG island of the TOX L1HS locus appeared densely methylated in all but one haplotype, which had half the baseline methyl-CpG score. Likewise, hypomethylation of UXS1-overlapping L1HS also appeared relatively rare, with two hypomethylation events separated by three generations in the Platinum pedigree. While variable promoter methylation over the XPR1 L1HS locus was present in our trio, this locus appeared hypermethylated throughout the Platinum Pedigree. By contrast, the two intergenic L1HS loci at chr10q25.1 and chr15q15.3, as well as the SCFD1 L1HS, showed multiple hypomethylation events in the larger pedigree. No clear Mendelian inheritance pattern was evident for SCFD1 or chr10q25.1, and not enough information was available to trace the inheritance of all hypomethylation events in chr15q15.3 due to missing data and insufficient coverage in some samples.

**Figure 4.**
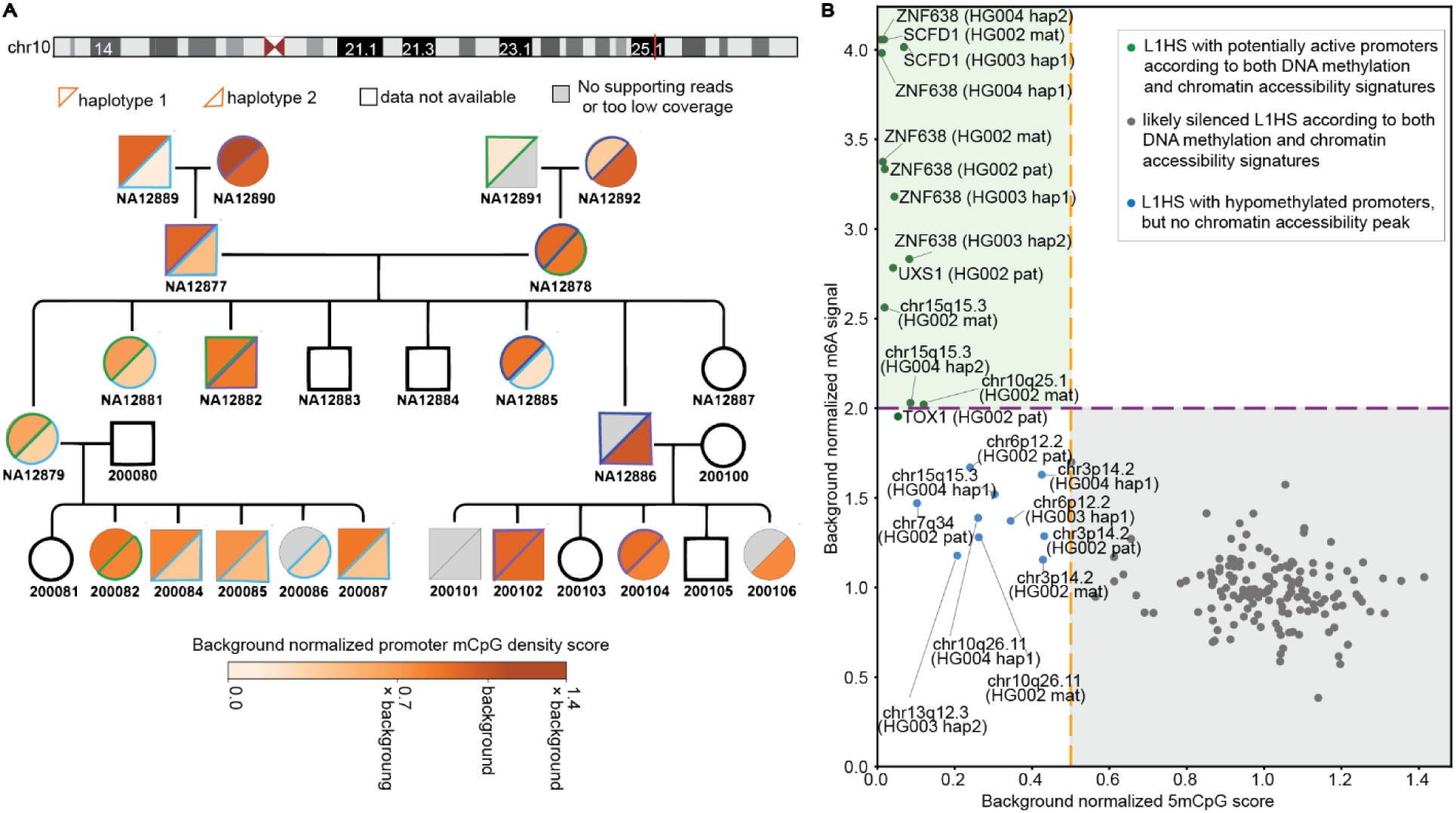
L1HS promoter methylation can vary discordantly with chromatin accessibility across generations. **A)** Platinum Pedigree view of haplotype-resolved promoter methylation at the chr10q25.1 L1HS locus. This locus shows variable methylation across generations, with the “green” and “blue” haplotypes showing signs of promoter methylation that does not strictly follow Mendelian inheritance. If the data for one or more haplotypes is missing, that is either because the reads over this particular L1HS could not be haplotagged, or the coverage of the region was fewer than 8 reads per haplotype. **B)** Comparison of background-normalized promoter 5mCpG density scores and m6A signals in trio samples over loci with hypomethylated promoters in the Platinum Pedigree. Dashed lines are visual references at half the mean 5mCpG score and twice the mean m6A score. Loci separate into three groups (colored dots): candidate escape loci with promoter hypomethylation (lower outlier 5mCpG score) and increased chromatin accessibility (upper outlier m6A signal) with FIRE enrichment, likely silenced L1HS with high promoter methylation and low accessibility, and hypomethylated loci with no FIRE enrichment and no outlier m6A signal, suggesting that promoter hypomethylation alone is not sufficient for L1HS activation.

Next, we investigated whether any additional near-full-length (>5.9kb) L1HS elements within the Platinum Pedigree^42^ showed evidence of promoter hypomethylation. We identified 10 additional L1HS loci with promoter methyl-CpG density scores less than half of the background level in multiple individuals, as well as 18 loci with similarly reduced scores in only a single individual and haplotype within the pedigree (Supplementary Table 2). To identify candidate non-Mendelian methylation patterns, we compared each offspring haplotype with the corresponding parental haplotype. We required both a discordant hypomethylated/not-hypomethylated label and a difference in background-normalized methylation greater than 20 percentage points, similar to the effect-size cutoff used in a previous human trio whole-genome bisulfite sequencing study^43^. We considered a locus informative only if it had sufficient coverage after phasing for at least two parent-offspring transmissions. By these criteria, half of the multi-individual loci and 8 of the 18 single-individual/haplotype loci showed methylation patterns inconsistent with simple co-inheritance of methylation state with the transmitted haplotype (Supplementary Figure 5 and Supplementary Table 3). Including the previously described L1HS loci with non-Mendelian inheritance in our trio data, and counting only autosomal L1HS loci, this corresponds to ∼6% of autosomal near-full-length informative L1HS loci showing potential non-Mendelian inheritance of promoter DNA methylation patterns (17/290). Although these percentages are more observational, as pedigree data does not have replicates needed for more robust statistical testing of methylation differences between generations, this frequency is similar in magnitude to the observation by Davidovich et al.^44^ that approximately 7% of identified autosomal epigenetic inheritance patterns in mice were non-Mendelian.

While some of the hypomethylated L1HS elements identified in the Platinum Pedigree, particularly those hypomethylated in only a single haplotype, may have truly escaped silencing, this is unlikely to explain all cases, as L1HS reactivation is thought to be rare in healthy differentiated cells. We therefore hypothesized that, although DNA methylation at these loci was variable, chromatin accessibility might remain low at least in some cases, providing an additional barrier to reactivation. This hypothesis is in line with Lanciano et al.’s^7^ observation that for over 80% of L1HS loci, even 5-aza-2′-deoxycytidine-induced DNA hypomethylation does not lead to detectable transcription from their promoters. In confirmation of this, we find that all hypomethylated L1HS loci identified in our analysis remain transcriptionally silent after 5-aza-2′-deoxycytidine treatment. Because the Platinum Pedigree dataset does not include chromatin accessibility information, we examined the same loci in our trio dataset to determine whether hypomethylation was consistently associated with increased accessibility. Aside from the loci described above that showed both promoter hypomethylation and accessible chromatin in at least one trio haplotype (ZNF638, SCFD1, UXS1, TOX, chr10q25.1, and chr15q15.3), the remaining L1HS loci hypomethylated in the Platinum Pedigree did not show evidence of accessible chromatin across trio haplotypes by FIRE peak-calling and outlier-threshold methods. For most of these loci, lack of accessible chromatin in the trio was accompanied by promoter hypermethylation. In contrast, five loci remained inaccessible despite showing promoter hypomethylation in at least one haplotype in the trio, a pattern also observed for one haplotype of the previously described chr15q15.3 L1HS (Figure 4B).

### Fiber-TEnCATS complements long-read-based assemblies and enables more accurate inheritance tracing of L1HS insertions

Moving away from L1HS loci present in all cell lines of our trio, we next focused on polymorphic L1HS insertions in the trio. Even though the parents’ personal patched genomes, HG003 and HG004, were generated from long-read-based assemblies (Methods), their representation of L1HS loci remained incomplete. With Fiber-TEnCATS, we identified 20 candidate near-full-length (>5.9 kb) parent-specific L1HS copies that were missed in the corresponding parental assembly, none of which were inherited by HG002 based on its T2T assembly. Fiber-TEnCATS also enabled inheritance tracing of the 15 near-full-length L1HS insertions heterozygous in the HG002 T2T assembly but absent from both parental assemblies (Figure 5A and Supplementary Table 4). As these copies were missing in the parents’ assemblies, these heterozygous L1HS in HG002 could not have been explained by inheritance and would falsely inflate the potential *de novo* L1HS count in HG002, especially since all except one of these have no frameshift mutations or premature stop codons in either of their ORFs (Supplementary Table 4).

**Figure 5.**
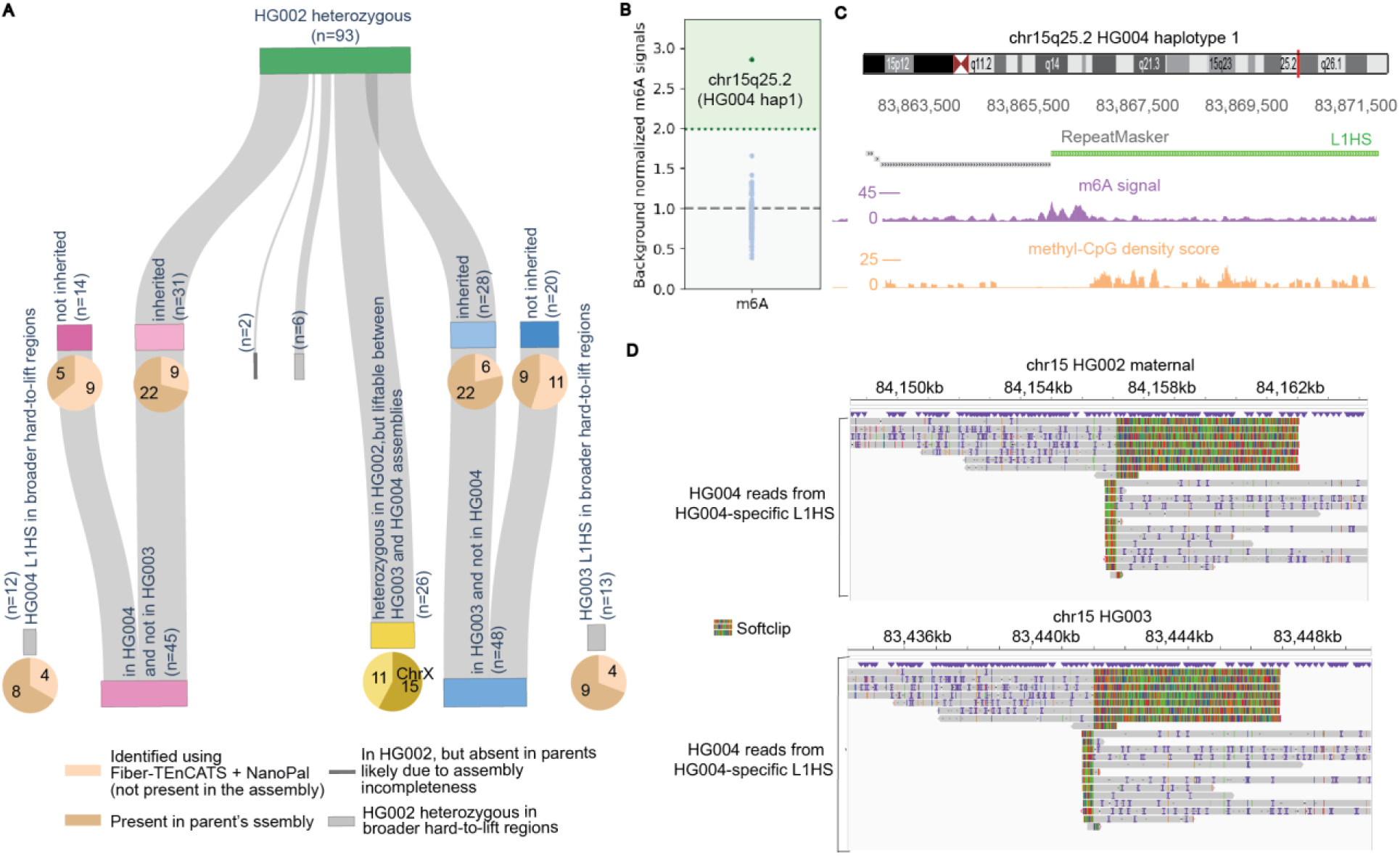
Chromatin accessibility over a polymorphic L1HS insertion on chr15 in HG004 that is absent from two other trio samples. **A)** Inheritance of (near) full-length L1HS that are absent in at least one of the cell lines in the trio. Because the parental assemblies are not haplotype-resolved, counts are shown per locus rather than per allele. **B)** Normalized promoter m6A signals for all polymorphic L1HS insertions from A). **C)** GBiB view of L1HS locus on chr15q25.2 with accessible promoter in HG004 **D)** IGV view of reads from L1HS locus on chr15q25.2 in HG004 aligned to HG002 and HG003 genomes. L1HS part of each read appears as a softclip, suggesting that this L1HS is absent from HG002 and HG003.

Among the total of 93 heterozygous HG002 L1HS insertion loci, we identified 2 loci (chr2q31.1-paternal and chr8q21.11-maternal) that could not be explained by inheritance from the augmented parents’ set of L1HS or by residing in the broader “hard-to-lift regions” (Methods). However, neither of these two L1HS elements is likely to be a true *de novo* insertion in HG002. The chr2q31.1 L1HS contains several premature stop codons, making it unlikely to be a newly active insertion. The chr8q21.11 L1HS has intact ORFs, with no detected frameshift or nonsense mutations, but is also unlikely to be *de novo* because an L1HS insertion at the same locus has been observed in several other samples not related to HG002^38,45,46^. The most probable explanation for why these two L1HS elements were not recovered by Fiber-TEnCATS, unlike the 15 previously mentioned L1HS elements that were similarly heterozygous in HG002 and absent from the parental assemblies, is that chr2q31.1 and chr8q21.11 have mutations in their target sites and are thus not efficiently recognized by the Cas9/sgRNA complex. The chr2q31.1 L1HS contains seven mismatches to our sgRNA, and the chr8q21.11 L1HS has a deletion of one G required in the canonical NGG PAM.

We proceeded to investigate promoter accessibility and methylation of all L1HS from Figure 5A and found that almost all of them had promoter m6A levels resembling the silenced L1HS signature, reflecting the host cell’s tendency to silence evolutionarily young TE copies (Figure 5B). However, we identified a single HG004-specific heterozygous intergenic L1HS insertion on chr15q25.2 with an accessible, hypomethylated promoter (Figures 5C and 5D), but no detectable transcripts in HG004 Iso-seq data. Insertion polymorphisms of this L1HS have been previously reported^38,46,47^, with progeny of its transposition observed in several cancer tissues^38,46^. While L1Xplorer ^48^ found multiple frameshift mutations in the HG004 personal genome-derived ORF2 sequence of this L1HS, the sequence reconstructed from Fiber-TEnCATS reads over this locus had intact open reading frames (Supplementary Table 5). Due to the opposing implications of these two sequence compositions for the potential transposition competence of this L1HS locus in HG004, we checked for other evidence of recent activity, such as preserved target-site duplications (TSDs) and likely sources or progenitors of this L1HS. We identified a 14bp-long TSD on either side of this L1HS locus, based on both genomic and read-derived sequences, and a transduction sequence (Supplementary Table 5) with an intact tag specific to the lineage of AC002980 ‘hot’ L1HS (“GCTTTATTGAGGTGTAACCAGCA”)^22^. Both the transduction sequence and chr15q25.2 L1HS internal sequence showed over 99% identity to the ‘hot’ chrXp22.2 L1HS locus^22,37,38,45–47,49^, present in HG004 and inherited by HG002 with both intact ORFs (Supplementary Table 5).

## Discussion

Overcoming the multimapping and limited haplotype resolution of short-read technologies while avoiding the high costs of long-read whole-genome sequencing methods can be a challenge. To address these limitations, we developed a TE-targeted approach that provides higher throughput at lower costs and enables simultaneous analysis of allele-specific sequences, DNA methylation, and chromatin accessibility across the full length of both reference and non-reference L1HS loci. With this approach, named Fiber-TEnCATS, we comprehensively characterized L1HS DNA methylation and chromatin accessibility in a human lymphoblastoid trio (HG002, HG003, and HG004).

Antisense transcripts of L1HS locus in an intron of ZNF638 (NP220) have been previously identified in several cell lines and tissues^4,8,10^, and promoter CpG hypomethylation has been observed in healthy heart tissue^30^. Here, we extend previous locus-level characterization by overlaying multiple haplotype-resolved markers of promoter activity on this L1HS across multiple family members and observe consistent methyl-CpG patterns in a larger pedigree. This observation led us to speculate that the active state of this L1HS’s promoter might be maintained by the host cell rather than resulting from sporadic escape from silencing. The position of this L1HS within the ZNF638 gene and the inclusion of ZNF638 exons in antisense transcripts made this possibility especially intriguing, as the HuSH-ZNF638 interaction plays a role in silencing extrachromosomal retroviral and AAV DNA^50,51^, while the ZNF638 interaction with TASOR could impact LINE silencing^52–54^. While further research is needed to address this hypothesis, we speculate that regulating the activity of this L1HS could be one of the host cell’s mechanisms for fine-tuning ZNF638 levels, thereby modulating the downstream effects of the ZNF638 interactome on the LINE silencing. Our sequence analysis indicates that maintaining the promoter of this L1HS active is unlikely to increase genomic instability by allowing this L1HS to retrotranspose autonomously, contribute to the movement of non-autonomous TEs, or increase the pool of ORF2p endonuclease activity, as ORF2 of this L1HS contains a premature stop codon.

Our study also highlights the significance of read haplotagging when evaluating the activity of genomic L1HS copies. Long reads and high coverage from Fiber-TEnCATS across L1HS loci revealed heterozygous epialleles in six L1HS. This variation in L1HS accessibility would have gone unnoticed from transcription data alone, as 2 of these L1HS don’t have detectable transcripts, and transcripts from 3 of the other 4 L1HS could not be assigned to a specific haplotype, despite being derived from long reads.

Recently, Davidovich et al.^44^ found that approximately 7% of identified autosomal DNA methylation inheritance patterns in mice deviated from Mendelian inheritance. Our promoter DNA methylation analysis of near-full-length L1HS loci in the trio data and the larger four-generation family suggested a similar magnitude, with ∼6% of informative loci showing patterns inconsistent with simple co-inheritance of methylation state with the transmitted haplotype.

However, we note that human pedigree analyses lack the biological replication of controlled mouse crosses and the statistical power of population-scale association studies and that, therefore, these results should be viewed as observational evidence for potential differences across generations, rather than as replicate-based tests of robust methylation inheritance. The absence of a clear Mendelian pattern in the DNA methylation inheritance at these L1HS loci makes it unlikely that their promoter hypomethylation was driven by *cis*-acting meQTL. However, a much larger sample size will be needed to determine whether and which *trans-*acting genetic variants influence the observed CpG methylation changes and their inheritance at specific L1HS loci. Hypomethylation of at least some L1HS loci, particularly those hypomethylated only in a single cell line in our analysis, might be a consequence of stochastic variability in transposon silencing rather than genetically driven effects. This would be consistent with the broader view that most epigenetic states are transient across generations unless stabilized by underlying genetic variation^55–57^. We also note that donor age may contribute to the observed epigenetic variability, as L1HS promoter methylation has been reported to decrease with age^58^, and that sample type may also contribute, since lymphoblastoid cell lines can acquire epigenetic alterations during EBV transformation, immortalization, and culture^59,60^.

Supporting prior locus-level work findings^7,24,25^ that hypomethylation alone is not sufficient to imply an open promoter state, our Fiber-TEnCATS trio data revealed that observed DNA methylation variability is not always reflected in L1HS chromatin accessibility. This underscores that L1HS regulation relies on multiple silencing layers and that simultaneous measurement of DNA methylation and chromatin accessibility improves our understanding of the L1HS regulatory state.

High sequencing depth across L1HS loci that Fiber-TEnCATS provides is important not just for epigenetic analysis but also enables the discovery of L1HS insertions that even personal long-read-based assemblies miss. With our method, we recovered 15 L1HS insertions that were absent from the HG003 and HG004 assemblies but were present in one parent and inherited as heterozygous insertions in HG002. Resolving these parental alleles allowed us to trace their inheritance and distinguish inherited insertions from loci that may have appeared consistent with *de novo* insertions in HG002 based on assembly comparison alone, especially because all but one lacked frameshift mutations or premature stop codons in either ORF. Fiber-TEnCATS also identified additional candidate parent-specific insertions missed by parent assemblies and not inherited by HG002.

Taken together, Fiber-TEnCATS is a cost-effective long-read approach for haplotype-resolved profiling of DNA methylation and chromatin accessibility at both reference and non-reference L1HS insertions.

## Methods

### Cell Culture

GM24385, GM24143 and GM24149 lymphoblastoid cell lines were acquired from the Coriell Institute for Medical Research and were grown in RPMI 1640 media (Invitrogen, 11875093) supplemented with non-heat inactivated fetal bovine serum (Corning, 35010CV) at 37 degrees Celsius with 5% CO2. Cell density and viability were determined using a Countess II FL (Invitrogen).

### Hia5 chromatin labeling and genomic DNA extraction

Low passage (5 or below) lymphoblastoid cells were harvested from culture and washed in 1x phosphate buffered saline (PBS, Invitrogen, 10010023). Washed cells were resuspended in 10 mL of 1x PBS and counted using a Countess II FL. For each genotype, 4 million cells were aliquoted into a 1.5 mL eppendorf tube per replicate. Aliquoted cells were centrifuged in a swing bucket rotor at 500xg for 5 minutes. After aspirating the supernatant, cell pellets were resuspended in 250 μL of Buffer A (15 mM Tris-HCl pH 8.0, 15 mM NaCl, 60 mM KCl, 1 mM EDTA pH 8.0, 0.5 mM EGTA pH 8.0, and 0.5 mM Spermidine). To isolate nuclei, 250 μL of 2x Lysis Buffer (Buffer A with 0.6% IGEPAL) was added to the cell aliquots resuspended in Buffer A and incubated on ice for 10 minutes. Nuclei suspensions were centrifuged at 350xg for 5 minutes in a swing bucket centrifuge and the supernatant aspirated. The nuclei pellet was resuspended in 232.5 μL of Buffer A with 6 μL of 32 mM S-Adenosylmethionine (SAM, NEB, B9003S) and 1.5 μL of Hia5 enzyme (200U/μL, provided by Andrew Stergachis) and incubated at 25°C for 10 minutes. After Hia5 labeling, nuclei were centrifuged at 350xg for 5 minutes in a swing bucket rotor and the supernatant aspirated. Hia5 labeled pellets were immediately prepared for genomic DNA (gDNA) extraction.

Genomic DNA of Hia5 labeled nuclear pellets was extracted using a modified salting out method ^61^. Pellets of 4 million Hia5 labeled nuclei were resuspended in 500 μL of lysis buffer (10 mM Tris HCl pH 8.0, 400 mM NaCl, 2mM EDTA pH 8.0). To the resuspended nuclei, 40 μL of 10% SDS and 80 μL of Proteinase K (10 mg/mL) were added and incubated at 55°C for at least one hour. Following the incubation, 200 μL of a solution of saturated NaCl was added and the lysate vigorously shaken to mix, followed by centrifugation at 4000xg for 15 minutes. The clarified supernatant was then poured into 2 volumes of 200 proof ethanol and gently inverted end over end for 5 minutes at room temperature (RT) using a tube rotator. The gDNA was spooled out using a p10 filter tip and re-suspended in 60 μL of TE buffer (10 mM Tris-HCl, pH 8.0, 1mM EDTA, pH 8.0) at 4°C overnight.

### Transposable element nanopore Cas9 targeted sequencing (TEnCATS)

TEnCATS was performed based on the protocol in Van Deynze et al.^62^ with the following changes briefly described here. Purified genomic DNA was quantified using a Qubit fluorometer (Invitrogen) and 30 μL (at a concentration of 100 ng/μL or greater) was dephosphorylated. The RNP was produced by combining 850ng of *in vitro* transcribed L1HS oligo used in McDonald et al. (2021), 1.5 μL 1X Cutsmart buffer (NEB, B7204S), 1 μL of a 1:5 dilution of Alt.-R *S.p.* HiFi Cas9 nuclease V3 (IDT, 1081058) to 1X Cutsmart Buffer, and dH2O to bring the reaction to a total of 15 μL then incubated at RT for 20 minutes. After dephosphorylation, the RNP was added to the gDNA followed by 1.5 μL of dATP and 2 μL of Taq DNA polymerase (NEB, M0273S) and incubated at 37°C for 30 minutes then 72°C for 10 minutes. 2.5 μL of thermolabile proteinase K (NEB, P8111S) was added following the Cas9 cutting-polyA-tailing and incubated at 37°C for 10 minutes then 72°C for 10 minutes to remove residual enzyme. The reaction went into the adapter ligation followed by addition of 0.4x Axygen magnetic beads (ONT, SQK-LSK114) and 2x washes with LFB (ONT, SQK-LSK114). The library was eluted in 21 μL of EB (ONT, SQK-LSK114) then incubated at 37°C for 10 minutes. The eluate was placed into a new 1.5 mL microcentrifuge tube and the concentration determined by Qubit. 20 μL of library was loaded onto a promethION R10.4.1 flow cell following the ONT protocol and sequenced for 72 hours with a single wash and reload at 24 hours using the same library and ONT’s protocol.

### Constructing personal genome references

To better represent the L1HS landscape in the trio cell lines and improve read mapping, we used personal genomes instead of the GRCh38 reference. For HG002, we used the Telomere-to-Telomere Consortium, Human Pangenome Reference Consortium, and the Genome in a Bottle Consortium’s (GIAB) HG002 T2T haplotype-resolved personal genome v1.0.1 (https://github.com/marbl/HG002)^63^. Since HG003 and HG004 do not have publicly available T2T genomes, we chose the closest alternative—personal patched genomes^64^ created by filling gaps in the publicly available GIAB PacBio long-read genome assemblies for HG003 (GCA_001549605.1) and HG004 (GCA_001549595.1) with the corresponding CHM13 T2T sequence using GPatch^63^.

### Transposable elements annotation and identification

RepeatMasker 4.1.6^65^ was used to obtain the initial pass of transposable element annotations for all genomes. RepeatMasker occasionally artificially truncates L1HS annotations, especially those inserted as reverse complements. This is because members of the LINE family do not have full-length consensus sequences in the database that RepeatMasker uses, so RepeatMasker scans for a highly conserved ORF2 sequence and consensus sequences for 5’ and 3’ UTRs of specific subfamilies separately and assembles them while adjusting all positions to the position of ORF2 in a complete L1PA2 element. To define L1HS boundaries more precisely, we ran BLAST^66^ on RepeatMasker-identified genomic L1HS sequences against the sequence of a full-length transpositionally active L1HS (L1.3 GenBank accession no. L19092)^67^ and adjusted the genomic coordinates and match to the L1HS sequence accordingly. To make the repertoire of L1HS in HG003 and HG004 more complete, we used NanoPal^29^ on Fiber-TEnCATS reads to identify L1HS insertions missed by patched personal genomes. From the full set of NanoPal identified non-reference L1HS, we kept only those with at least 8 supporting reads and longer than 5.9kb. This length filter reduces the likelihood of false-positive calls because errors in long-read assemblies can generate soft-clipped alignments that NanoPal may misinterpret as short non-reference insertions (Supplementary Figure 6). We also developed a script to reconstruct long NanoPal L1HS sequences from their supporting reads, with an option to either incorporate them into the assembly or write them to a VCF file with the inferred insertion position and corresponding sequence. For each candidate insertion, sequences were reconstructed independently from reads spanning each side of the Cas9 cut site. In the soft clip of each read supporting the insertion, we identified L1.3-matching sequence nearest to the cut site using BLAST, grouped reads supporting the same breakpoint, aligned the corresponding soft-clipped sequences with MAFFT^68^, removed alignment columns with gaps in more than half of supporting reads (if any), and generated consensus sequences using EMBOSS^69^.

Individual-specific genomic copies of L1HS, or haplotype-specific in the case of HG002, were identified using the UCSC LiftOver tool^70^ to map L1HS positions between different genomic coordinate systems. Chain files between personal genomes were created using the T2T consortium pipeline available at the UCSC website: https://genome.ucsc.edu/cgi-bin/hgTrackUi?hgsid=1362445199_8KVk8xT77Y7RKwC4d3AbkJb8E4ZL&db=hub_3267197_GCA_009914755.4&c=CP068275.2&g=hub_3267197_hgLiftOver. To identify L1HS insertion events present in only a subset of the trio cell lines, we restricted the analysis to near full-length L1HS elements, defined as insertions with less than 100 bp of 5′ truncation and a total length greater than 5.9 kb. Near-full-length L1HS insertions from each genome were retained as candidate variable insertions if they failed to lift over to the genomic coordinate system of at least one other trio member. Insertions originating from CHM13 patches in HG003 and HG004 patched personal genomes were excluded from the parent-specific set. Inheritance of NanoPal-identified L1HS insertions was additionally verified by lifting over 5 kb upstream and 5 kb downstream flanking sequences from each parental genome to HG002 and then determining whether an L1HS insertion was present between the mapped flanks. All homozygous L1HS insertions in HG002 were classified as present in both parents, regardless of whether they were explicitly represented in the parental assemblies. Insertions that could not be evaluated because they fell within broader hard-to-lift regions were assigned to a separate category. A hard-to-lift region was defined as one in which less than 1% of the 20 kb sequence upstream or downstream of the L1HS insertion could be lifted over.

### L1HS source inference

L1HS source inference based on the internal L1HS sequence was performed as in Wang et al.^39^ – after trimming poly-A tails, the selected full-length L1HS was compared to all other full-length L1HS using BLAST, and the L1HS with the highest unique BLAST score was deemed a potential progenitor or progeny. In parallel, L1HS source loci were inferred using transduction-sequence similarity. For L1HS loci with an additional poly-A tail within 2 kb of the 3′ end of the internal L1HS sequence, the intervening sequence was compared to the 2 kb flanks of all other full-length reference and non-reference L1HS loci using BLAST. Intervening sequences longer than 50 bases that have ≥85% identity to another locus were classified as candidate transductions, and the highest-bitscore match was assigned as the putative source or progeny locus. L1Xplorer^48^ was used to identify any frameshift or nonsense mutations in ORFs of L1Hs.

### Assessing chromatin accessibility and DNA methylation

DNA basecalling was performed using Dorado (https://github.com/nanoporetech/dorado) with the dna_r10.4.1_e8.2_400bps_sup@v5.0.0 simplex basecalling model and the 5mCG_5hmCG and 6mA modified-base models. At the read level, a likelihood cutoff of ML>225 was applied to assign methylation status to bases, in accordance with Jha et al.^71^, who demonstrated that at this threshold, the percentage of Dorado-called m6A sites best approximates the m6A levels quantified via MS/MSl. The unaligned BAM files were converted to FASTQ format, and methylation tags were retained using samtools^72^ fastq -T "*". Reads were aligned to the appropriate personal genome assemblies using minimap2^73^ with the following parameters: -ax map-ont --eqx -Y -y --secondary=no -t 20. Read haplotagging was performed as described in the "Reads haplotagging" section. For each adenine and CpG position within L1HS, methylation percentage was calculated as the proportion of reads classified as methylated at that position. Because L1HS silencing depends on CpG methylation as well as local CpG density, we quantified CpG methylation using a methyl-CpG density score, defined as the sum of methylation probabilities across all CpG sites within a window divided by the total number of cytosines in that window, or guanines for reverse-strand reads. m6A signal was calculated analogously as the sum of methylation probabilities across all adenines within a window divided by the total number of adenines, or thymines for reverse-strand reads. To visualize chromatin accessibility and DNA methylation along the full L1HS, m6A signal and methyl-CpG density score were averaged across 100-bp sliding windows and converted to BigWig format with kentUtils (https://github.com/ENCODE-DCC/kentUtils).

Aligned BAM files were processed with Fibertools’ –track-decorators^71^ to generate BED and decorator files, which were subsequently converted to .bb format using kentUtils for visualization in the Genome Browser in a Box^74^. Deeptools^75^ computeMatrix scale-regions and plotHeatmap were used to create m6A metagene plots over ATAC-seq regions, all scaled to the same length. For this, HG002 reads were aligned to the GRCh38 reference genome, and GM12878 ATAC-seq (ENCSR637XSC) peak coordinates were obtained from the ENCODE database (https://www.encodeproject.org/experiments/ENCSR637XSC/).

L1HS promoter accessibility was assessed by calling Fiber-seq Inferred Regulatory Elements (FIREs) on individual reads using fibertools^34,71^, followed by capture-aware promoter-level FIRE enrichment testing. The original FIRE FDR procedure was designed for genome-wide data, where real and shuffled FIRE scores can be compared across the full assayed genome to obtain a global score threshold. Because our coverage was restricted to the region around L1HS loci and not representative of the genome-wide background, we used a capture-aware empirical null instead. We performed 10,000 shuffles of capture reads over L1HS elements with at least 8 reads of coverage, preserving coverage at each L1HS promoter and adjusting read orientation so that parts of the read that overlapped promoters before shuffling, did so after shuffling as well. We then compared the observed mean FIRE score for each promoter to the distribution of mean scores from 10,000 random shuffles and applied the Benjamini–Hochberg correction to adjust for multiple testing. For this analysis, we defined a broader promoter region as the first 600 bases, in accordance with sense and antisense promoter positions defined in the literature^9,76^.

We used an independent promoter-level m6A outlier analysis to validate this modified FIRE-based approach. We normalized average m6A accessibility across the first 600 bp of each full-length L1HS promoter by dividing it by the average signal across all full-length L1HS promoters, which served as a silenced baseline. Because most full-length L1HS elements are expected to be silenced in healthy differentiated cells, we reasoned that active promoters should appear as outliers (Q3 + 3×IQR) with increased m6A accessibility signal. Using an equivalent outlier framework, we identified hypomethylated promoters as those with methyl-CpG density of the promoter CpG island located in the first 450 bp of the element^8^ (Supplementary Figure 1B) below Q1 − 3×IQR.

### Platinum Pedigree inheritance tracing

We used the Porubsky et al.^35^ long-read–based VCF, which contains merged variants across all samples (unphased), and assembly-based phased VCFs for each sample individually to generate a consensus set of merged, phased variants. This was done by retaining only assembly-based variants that were also present in the long-read VCF. This consensus variant set was then used to haplotag the reads for each sample in the Platinum Pedigree^35^. To determine which parental haplotype is being passed down to the offspring, we made a custom script that counts shared informative heterozygous alleles between each parental haplotype and each child’s haplotype at each L1HS locus. Transmission was first assessed within a 20 kb window centered on the L1HS element, and if not enough informative variants were found, a 100 kb window was used as a fallback. The parental haplotype with the greatest number of allele matches to the child was assigned as the transmitted haplotype.

To assess changes in L1HS promoter methylation across generations, promoter methyl-CpG scores were first normalized to the average across all full-length L1HS promoters in the same sample. Loci with normalized promoter methylation scores <0.5 were labeled as candidate hypomethylated loci rather than using an outlier-based threshold, which is more sensitive to read-coverage variation in lower-coverage whole-genome haplotype-resolved data. The final pool of near-full-length informative L1HS loci was defined as loci longer than 5.9 kb with at least eight reads of coverage after phasing for the transmitted haplotype in two or more parent–offspring transmissions. Candidate non-Mendelian methylation patterns were defined using criteria of >20-percentage-point difference in normalized methylation, based on the cutoff used in a previous human trio study^43^, and discordant hypomethylated/not-hypomethylated labels between transmitted parent-offspring haplotypes.

### L1HS transcription analysis

L1HS transcripts in HG002 and HG004 were identified from 2 publicly available PacBio Iso-seq datasets per cell line: 1) https://ftp-trace.ncbi.nlm.nih.gov/ReferenceSamples/giab/data_RNAseq/AshkenazimTrio/HG002_NA24385_son/Baylor_PacBio/ and https://ftp-trace.ncbi.nlm.nih.gov/ReferenceSamples/giab/data_RNAseq/AshkenazimTrio/HG002_NA24385_son/Google_PacBio/ for HG002 and 2) https://ftp-trace.ncbi.nlm.nih.gov/ReferenceSamples/giab/data_RNAseq/AshkenazimTrio/HG004_NA24143_mother/Baylor_PacBio/ and https://ftp-trace.ncbi.nlm.nih.gov/ReferenceSamples/giab/data_RNAseq/AshkenazimTrio/HG004_NA24143_mother/Google_PacBio/ for HG004. After adapter trimming with limma (https://github.com/PacificBiosciences/pbbioconda), reads were haplotagged (see Reads haplotagging) and aligned to personal genomes using minimap2^73^ with the following parameters: -ax splice:hq -uf --eqx -Y -y --secondary=no. FLAIR 2.1.0^77^ was used to correct misaligned splice sites in Iso-seq data using genome annotations and short-read identified splice junctions. We required a minimum mapping quality of 20 for all Iso-seq transcripts. Short-read-supported splice junctions were identified with STAR^78^ using poly-A selected RNA-seq data from the same cell lines: https://ftp-trace.ncbi.nlm.nih.gov/ReferenceSamples/giab/data_RNAseq/AshkenazimTrio/HG002_NA24385_son/UNC_Illumina/reads/ for HG002 and https://ftp-trace.ncbi.nlm.nih.gov/ReferenceSamples/giab/data_RNAseq/AshkenazimTrio/HG004_NA24143_mother/UNC_Illumina/ for HG004.

### Reads haplotagging

HG002 Fiber-TEnCATS reads were haplotagged with HaplotagLR^36^ using GIAB phased VCF v4.2.1 (Data and Code availability) after aligning to GRCh38. Haplotagged reads were then realigned to the respective haplotype of the HG002 T2T genome. DeepVariant v1.9.0^79^ was used to call variants from HG003 and HG004 Fiber-TEnCATS reads in the respective personal genome coordinate system, and UCSC Margin v2.3 (https://github.com/UCSC-nanopore-cgl/margin) was used to phase the obtained vcf before read haplotagging. Iso-Seq reads were haplotagged using WhatsHap v0.18^80^, using the same VCF files used for Fiber-TEnCATS reads for haplotagging.

## Data availability

Generated Fiber-TEnCATS trio datasets will be available in the NCBI Sequence Read Archive (SRA) as BioProject PRJNA1480362 upon publication. Patched personal genomes for HG003 and HG004 will also be publicly available on Zenodo as DOI: 10.5281/zenodo.20798533.

The "Telomere-to-Telomere" (T2T) Consortium’s HG002 reference v1.0.1 genome is available on the consortium’s GitHub repository: https://github.com/marbl/hg002. GenBank accessions for Genome in a Bottle (GIAB) Consortium’s long-read-based HG003 and HG004 assemblies used for the making of personal patched genomes are GCA_001549605.1 and GCA_001549595.1. GIAB’s Iso-seq and Illumina HG002 and HG004 expression data are available at the following FTP: https://ftp-trace.ncbi.nlm.nih.gov/ReferenceSamples/giab/data_RNAseq/AshkenazimTrio/ and their v4.2.1 phased VCF for HG002 was obtained from https://ftp-trace.ncbi.nlm.nih.gov/giab/ftp/release/AshkenazimTrio/HG002_NA24385_son/NISTv4.2.1/GRCh38/SupplementaryFiles/HG002_GRCh38_1_22_v4.2.1_benchmark_phased_MHCassembly_StrandSeqANDTrio.vcf.gz. Instructions on how to access Platinum pedigree Amazon Open Data are on consortiums GitHub: https://github.com/Platinum-Pedigree-Consortium/Platinum-Pedigree-Datasets. The GM12878 ATAC-seq data used in this study is publicly available at the ENCODE portal under the ENCSR637XSC accession.

## Code availability

Analysis code and NanoMEI code are available on GitHub at the following links: https://github.com/Boyle-Lab/Fiber-TEnCATS-manuscript and https://github.com/Boyle-Lab/NanoMEI

## Acknowledgements

We thank Andrew B Stergachis for sharing the Hia5 enzyme. We also thank all members of the Boyle lab for helpful discussions and feedback provided throughout the manuscript preparation process.

## Authors’ contributions

K.P. and A.P.B. conceived the project. K.P. designed and performed computational analysis, figure generation, and interpreted results with guidance from A.P.B. T.M. carried out the Fiber-TEnCATS wet-lab experiments. J.S. established and cultured cell lines and isolated gDNA. A.D. generated the HG003 and HG004 patched genomes. K.P. and A.P.B wrote the manuscript with input from all authors. A.P.B secured funding and supervised the work.

