## Supplemental Figures for "Fiber-TEnCATS reveals haplotype-specific chromatin accessibility and DNA methylation at human L1HS loci"

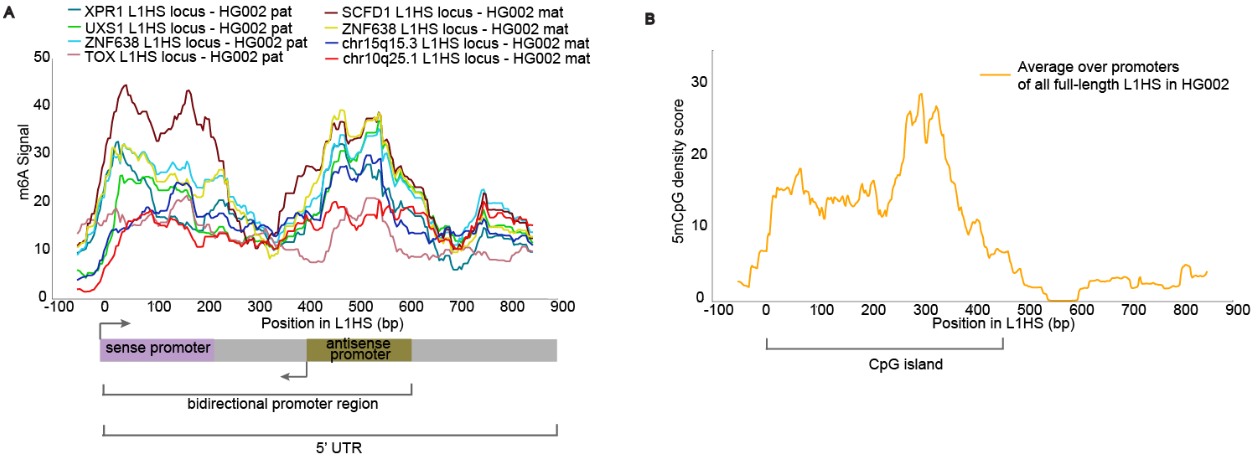


**Supplementary Figure 1. Locations of chromatin accessibility and DNA methylation peaks in L1HS 5'UTR** A) Raw m6A signal from open L1HS promoters, shown relative to the L1HS 5′ end. The two peaks line up with literature-reported coordinates for broader sense and antisense promoter regions B) Mean methyl-CpG density score across full-length L1HS promoters, shown relative to the L1HS 5′ end. The strongest signal is concentrated within the first ~450 bp of the promoter, overlapping the promoter's main CpG island. Since most L1HS loci are expected to be silenced and hypermethylated in differentiated cells, the average signal across all L1HS promoters provides an approximate silenced background.

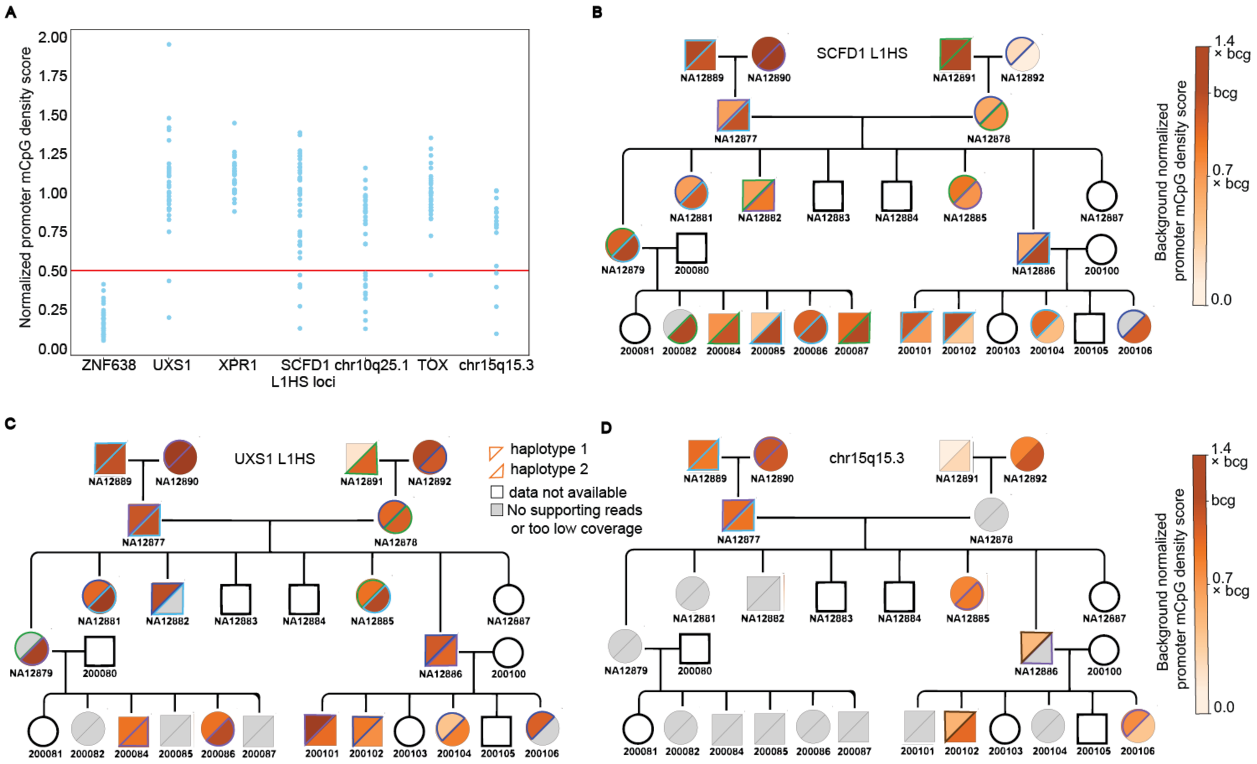


**Supplementary Figure 2. L1HS promoter methyl-CpG signatures in Platinum Pedigree** A) Average methyl-CpG density score across the first 450 bp of reference L1HS promoters with accessible promoter signal in the HG002 trio dataset, shown for haplotype-resolved samples from the Platinum Pedigree. For each sample, methyl-CpG density scores were normalized to the average score across all full-length L1HS promoters, which serves as an approximate hypermethylated/silenced background. Normalized scores lower than half of the expected silenced L1HS background were selected as candidate hypomethylated promoters. Panels B–D show a more detailed pedigree representation of promoter methyl-CpG density scores for three L1HS loci with multiple hyper and hypo methylation events in A (SCFD1, UCX1, and chr25q15.3).

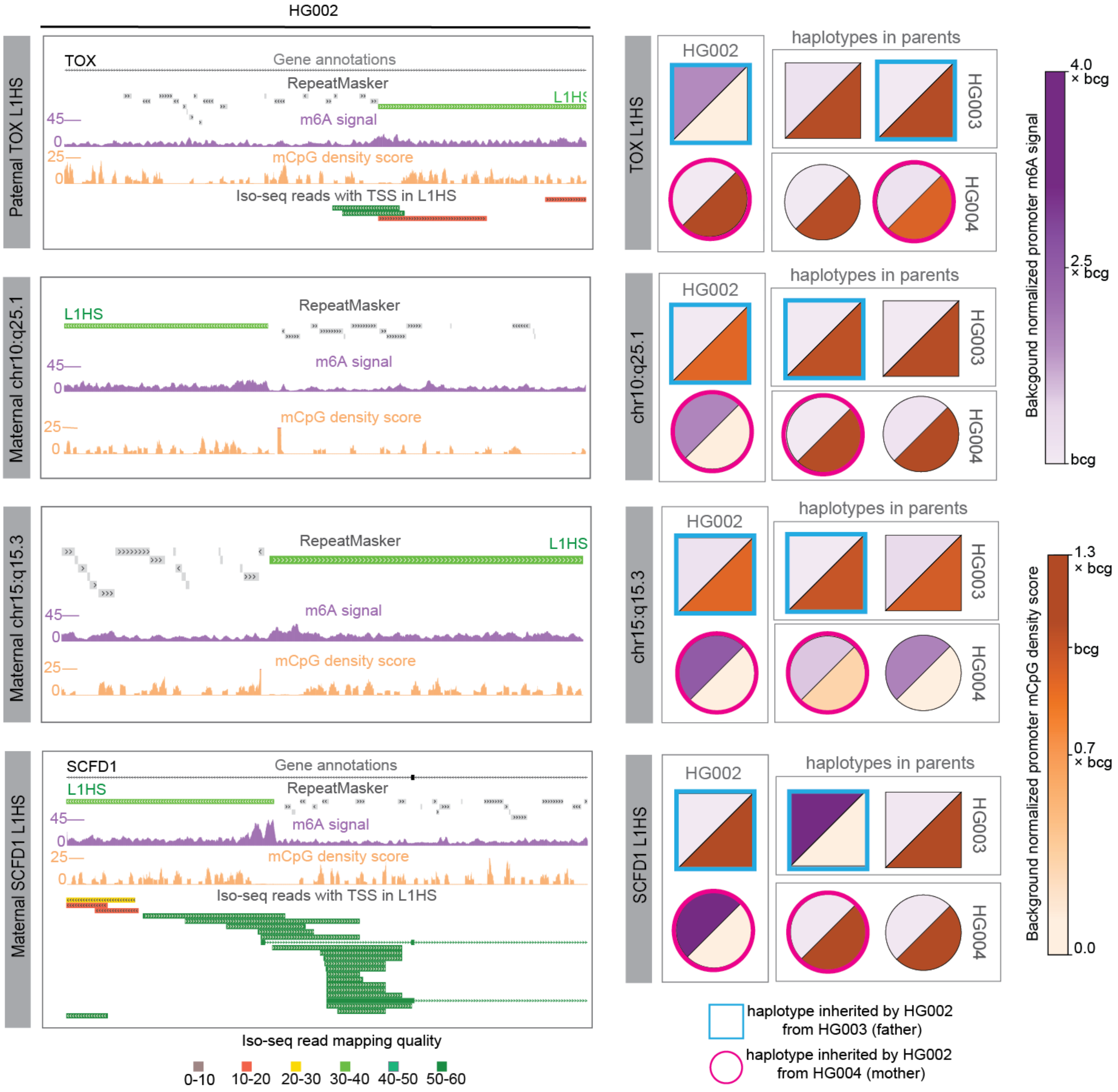


**Supplementary Figure 3. Haplotype-resolved view of TOX, chr10q25.1, chr15q15.3, and SCFD1 L1HS loci with heterozygous epialleles in HG002.** GBiB view of L1HS in the haplotype with detected m6A peak in HG002 (left) with corresponding haplotype-resolved and normalized DNA methylation (m5CpG) and chromatin accessibility (m6A) signals over promoters of L1HS in all trio samples (right).

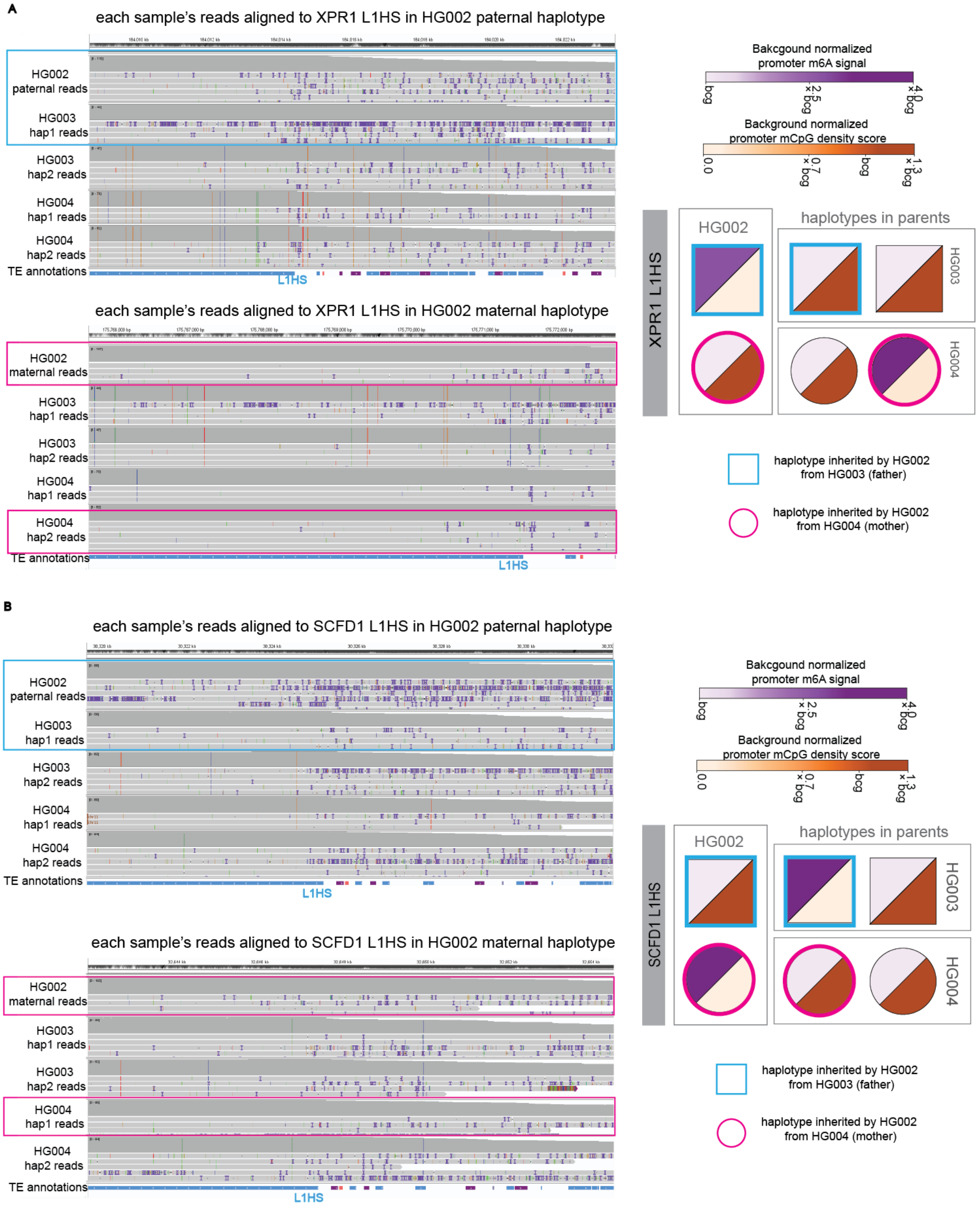


**Supplementary Figure 4. An IGV view of the XPR1 and SCFD1 L1HS locus showing informative variants supporting the assignment of inherited haplotypes.**

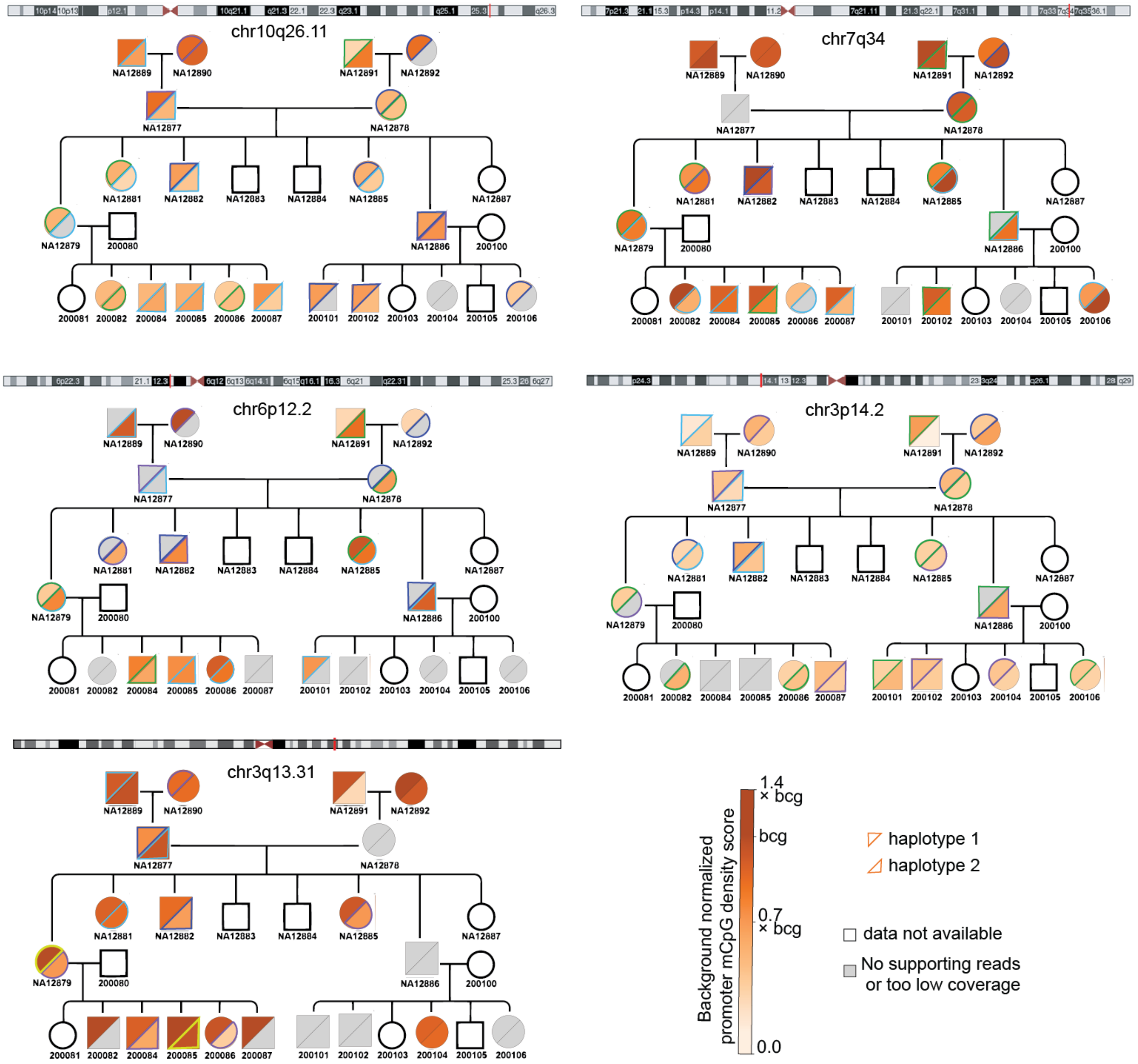


**Supplementary Figure 5. Examples of L1HS loci with variable promoter methyl-CpG density scores in Platinum Pedigree that do not follow Mendelian inheritance.** Scores were normalized to the sample-specific L1HS background, defined as the average methyl-CpG density score across all full-length L1HS promoters in each sample. Lower normalized scores indicate reduced promoter methylation relative to the expected silenced L1HS background. Missing values correspond to samples with either unavailable public data or haplotypes with less than 8 reads of coverage.

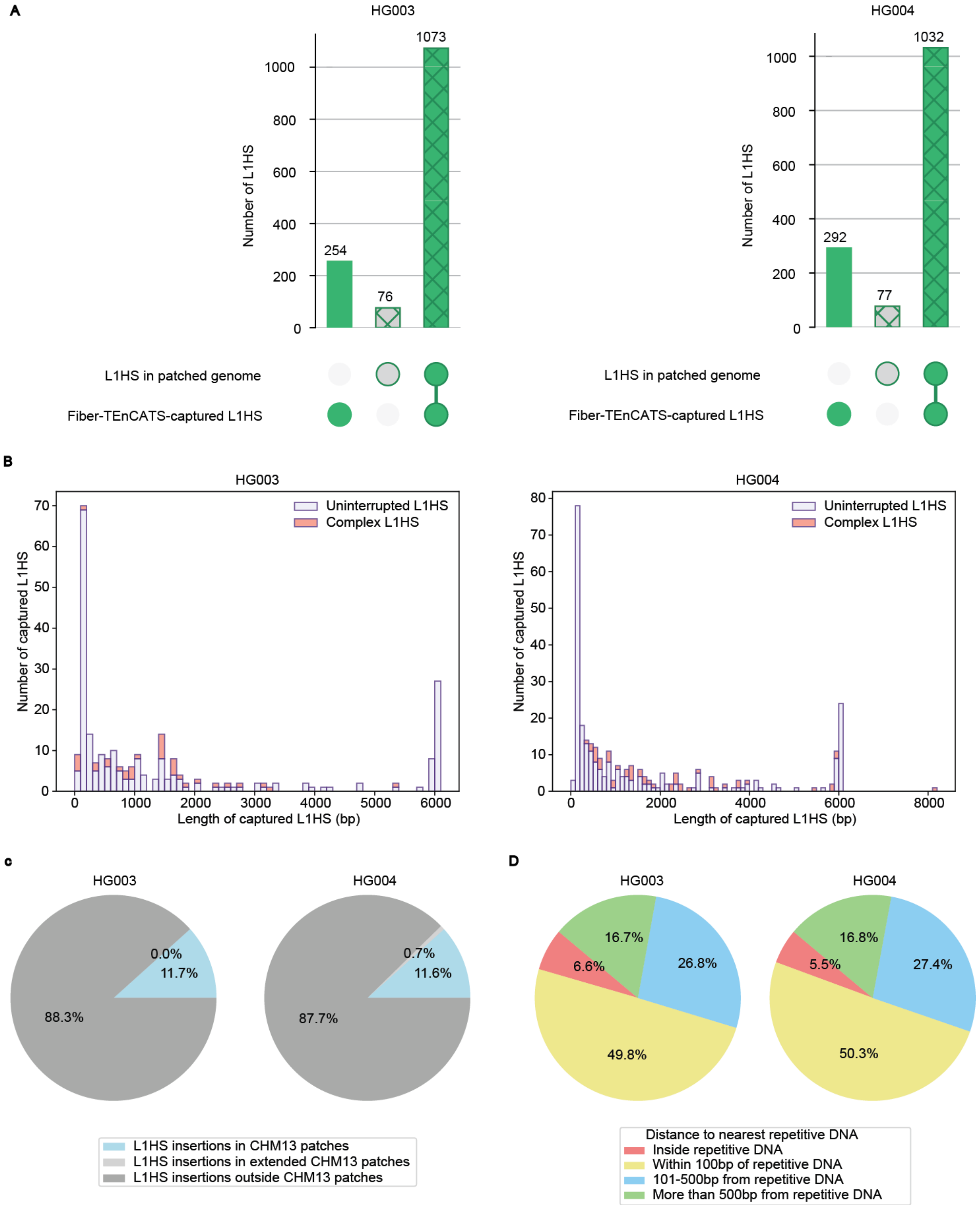


**Supplementary Figure 6: Summary of L1HS copies captured with Fiber-TEnCATS but absent from HG003 and HG004 personal patched genomes.** A) Summary of L1HS copies present in personal genomes versus those captured by Fiber-TEnCATS. The number of Fiber-TEnCATS-only insertions is likely inflated by short L1HS sequences appearing in read softclips due to assembly errors, rather than polymorphic insertions B) Length distribution of L1HS sequences captured by Fiber-TEnCATS but absent from the corresponding personal genomes. Overlapping or consecutive L1HS sequences within the same supporting-read softclip were classified as complex L1HS, with their combined length shown. C–D) Genomic context of missed L1HS sequences relative to genome patches (C) and repetitive elements (D), showing that our genome patching process is not the source of assembly errors inflating the count of NanoPal-identified non-reference L1HS insertions, as fewer than 1% of missed L1HS were located near patch boundaries. Most of NanoPal-identified non-reference L1HS insertions were found within 100 bp of repetitive regions that were likely difficult to assemble in the original genomes.
